# Functional MRI reveals regional changes of brain activity after five days of focal high-density theta burst stimulation (hdTBS) of the rat brain

**DOI:** 10.1101/2024.09.16.612091

**Authors:** Charlotte Qiong Li, Samantha Hoffman, Hieu Nguyen, Aidan Carney, Ying Duan, Zilu Ma, Nanyin Zhang, Yihong Yang, Hanbing Lu

## Abstract

**Background:** The therapeutic effects of transcranial magnetic stimulation (TMS) likely stem from neuroplasticity induced by repeated sessions over time. While animal models offer insights into TMS-induced plasticity, a rodent model that faithfully replicates prolonged TMS conditions in humans is still lacking.

**Objective/Hypothesis:** Develop a rat model that mimics the spatial and temporal patterns of TMS in humans.

**Methods:** Experiments were conducted on two cohorts of healthy adult rats (N=33). In cohort 1, rats underwent surgical implantation of microelectrodes for motor evoked potential (MEP) recording. With a rodent-specific coil and the high-density theta burst stimulation (hdTBS) paradigm, under awake condition, rats received daily TMS at 100% motor threshold for five days (days 1-5) to the hindlimb motor cortex. Cortical excitability was measured by input-output (I-O) curves on Day 0 (pre-hdTBS baseline) and Day 6 (post-hdTBS). The second cohort received identical TMS and underwent fMRI to map cerebral blood volume (CBV) on Days 0 and 6.

**Results:** Daily hdTBS session for 5 days significantly up-shifted I-O curves only in the TMS group (N=9), not in the sham group (N=7), indicating enhanced cortical excitability. fMRI data showed that, compared to sham group (N=9), rats receiving hdTBS (N=8) had increased basal CBV in several brain regions proximal and distal to the stimulation site, suggesting enhanced basal metabolism.

**Conclusion(s):** Daily hdTBS session for 5 days focally delivered to the motor cortex of naïve rats significantly altered basal brain activity in a network of brain regions, opening a novel platform for further investigating TMS-induced plasticity.

## Introduction

Therapeutic efficacy and time efficiency are two important factors in clinical application of transcranial magnetic stimulation (TMS). High-frequency repetitive TMS (rTMS) at 10 Hz and intermittent theta burst stimulation (iTBS) are the two most widely used protocols clinically. iTBS administers 600 pulses over 190 seconds, as opposed to 3000 pulses over 37.5 minutes in 10 Hz rTMS[1]. Both paradigms achieve similar, although modest, efficacy in treatment-resistant major depression [2–5]. The Stanford Neuromodulation Therapy (SNT) paradigm[6], an “accelerated” version of iTBS, achieved a higher remission rate. SNT applies 10 TMS sessions per day, each session delivers 1800 iTBS pulses with an inter-session interval of 50 minutes. Note that conventional iTBS was the building block of the SNT paradigm, which has been shown to exhibit limited effect in modulating cortical excitability[7]. There is a need to develop new TBS technology to enhance the efficacy while maintaining high time-efficiency, and to support further treatment optimization. For example, replacing the iTBS sessions in SNT with more effective TBS paradigms could potentially reduce the number of sessions required in a day while achieving high efficacy.

The therapeutic effects of TMS likely result from neuroplasticity that occurs over multiple sessions of TMS administration. For ethical reasons, testing of a new paradigm in healthy participants is usually limited to evaluating the after-effects in the motor cortex[1,8]. Therapeutic potential of the new paradigm is then assessed in patients, largely on a trial-and-error basis. A more systematic approach might involve developing TMS technology and assessing its acute and longitudinal effects in animal models first, with the aim of translating those results to human applications.

Back-translating TMS from humans to animals faces several technical challenges, with coil focality being a primary concern[9]. Small laboratory animals such as rats and mice require a small TMS coil to minimize off-target effects[10]; however, a small coil has a low efficiency [10,11], necessitating exceedingly high electric current to induce a suprathreshold response in the brain -- This leads to additional challenges such as coil overheating and electromagnetic stress. Recently, we developed a rodent-specific TMS coil with a focality of 2 mm[12,13]; Additionally, we introduced a high-density TBS (hdTBS) paradigm that administers up to 6 pulses per burst, compared to the 3 pulses in conventional TBS[14]. We assessed the acute after-effects of hdTBS based on motor evoked potential (MEP) readouts from awake rats, finding that 6-pulse hdTBS enhanced the after-effects by 92% compared to conventional 3-pulse iTBS.

Since longitudinal TMS, rather than acute TMS, holds greater therapeutic potential, in the present study, we aimed to determine whether longitudinal hdTBS sessions can induce neuroplasticity in healthy adult rats. Our previous observations indicated that an acute hdTBS session induced after-effects in healthy adult rats that last for about 35 minutes before returning to the pre-TMS baseline, suggesting that the brain circuits were temporally tipped “off-balanced”, and returned to balanced operation afterwards. Assuming that neural circuits in healthy brains operate at optimal efficiency as a result of evolution, and thus might be resistant to neuromodulation, the demonstration of neuroplasticity in healthy rats through longitudinal TMS would not only establish a model to investigate the mechanisms of TMS action, but also creates a novel and potentially significant platform for developing more effective TMS paradigms for human applications.

To this end, leveraging advancements in focal TMS technology[12] and awake rat TMS model[13], we administered daily single-session hdTBS to the motor cortex of awake rats for five days. We assessed MEP readouts before and after hdTBS modulation and mapped basal cerebral blood volume (CBV) using magnetic resonance imaging (MRI). CBV mapping, based on an exogenous MRI contrast agent, offers high resolution, high sensitivity, and is quantitative. Basal CBV is strongly coupled to basal cerebral blood flow (CBF)[15] and cerebral metabolism[16], reflecting brain activity. Our data reveal that one hdTBS session per day for five days significantly enhanced the excitability of the rat motor cortex, as measured by the input-output (IO) curve; CBV mapping showed that several brain regions, both proximal and distal to the stimulation sites, exhibited heightened basal CBV, suggesting increased regional brain activity. To the best of our knowledge, this is the first study demonstrating neuroplasticity induced by longitudinal, focal TMS of the motor cortex in awake rats. Our study opens a novel avenue for further investigating TMS mechanisms.

## Materials and Methods

### Animal preparation

Data were collected from 33 adult male Sprague–Dawley rats (14 to 16 weeks old) obtained from Charles River Labs. Of these, 16 rats were used for the MEP study (n=9 for hdTBS and n=7 for sham stimulation) and 17 for CBV mapping (n=8 for hdTBS and n=9 for sham stimulation). Surgical preparations are described below. Following surgery, the rats were individually housed under a reversed 12:12 hour light/dark cycle, with ad libitum access to standard rat chow and water. All experimental procedures were conducted following the guidelines established by the Animal Care and Use Committee of the NIDA-IRP and complied with the regulations outlined in the Guide for the Care and Use of Laboratory Animals.

### Microelectrode implantation for longitudinal MEP recording

MEP has been traditionally employed as a measure to assess the effects of TMS on the motor cortex[17,18]. In humans, the MEP signal can be conveniently recorded by measuring the electromyographic (EMG) signal from innervated muscles with surface electrodes. However, this method cannot be readily adapted to awake rats due to strong motion artifacts. We have developed a method that permits consistent MEP recording, detailed previously[14], and are illustrated in Figure 1A. Briefly, we customized EMG microwire-electrodes in house. The electrodes were made of soft 7-strand stainless steel microwires (0.025 mm in diameter, A-M systems, Washington, USA, cat. No. 793200), cut into13 cm in length. The insulation coating on one end of the microwires was removed (about 3 mm in length) before being press-connected to a female socket (Model E363/0, P1 Technology, USA). The sockets were then organized and embedded into a 6-channel electrode pedestal (Model: MS363, P1 Technology, USA). To ensure the stability of the sockets and microwires, the pedestal was attached to a circular Marlex mesh secured with dental cement. We custom-made a steel dust cap to cover the pedestal, which helped prevented the animals from chewing the plastic connectors, elongating the lifetime of the instrument during repeated TMS recording sessions. The insulation coating at the other end of microwires was also carefully removed, which were implanted into the biceps femoris muscle and the gastrocnemius muscle of the left hindlimb. These exposed segments served as active electrodes to detect EMG signal in the implanted muscles; one surface EEG pad was attached to rat tail serving as the ground electrode. The two active electrodes and the ground electrode were interfaced to a BIOPAC EMG recording system (BIOPAC Systems Inc, CA, USA). The EMG signal was amplified by a factor of 2000, band-pass filtered between 100 and 5000 Hz, and sampled at 10,000 Hz.

**Figure 1.**
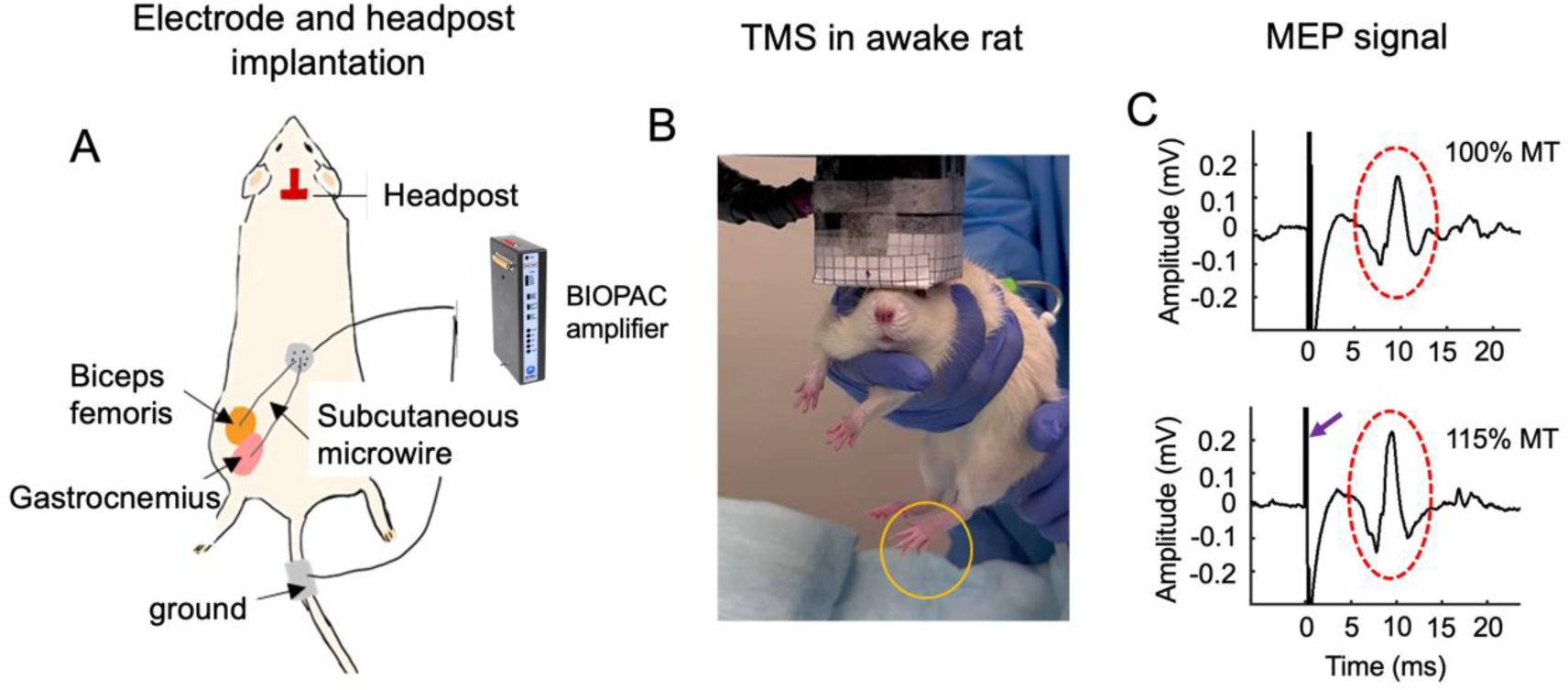
Illustration of experimental setup. (A): Customized microwire electrodes were surgically implanted into the biceps femoris and gastrocnemius muscles of the left hindlimb for longitudinal MEP recording; a headpost was implanted on the rat skull, serving as the reference for consistent coil positioning across TMS sessions and across animals. (B): An awake rat undergoing TMS administration. TMS pulse innervated the left limb (orange circle). (C) shows raw MEP traces at two power levels. The arrow indicates artifacts induced by a TMS pulse; dashed oval circles indicate MEP signal.

### Headpost implantation for consistent coil placement

A homemade TMS coil with a focality of 2 mm was used in this study. The high focality necessitates a method to consistently direct the hotspot of the TMS coil to the targeted brain region (hindlimb motor cortex). We implanted a headpost on the rat skull, which served as the reference to facilitate accurate coil positioning (see Figure 1A). Detailed procedures for headpost implantation were described previously[13]. Briefly, the primary motor cortex of the rat hindlimb has been mapped using classical electrical micro-stimulation (bregma coordinates: -1.8 mm anterior-posteriorly, 2 mm medio-laterally)[19]. A 3D-printed, T-shaped headpost was carefully implanted onto the rat skull such that the end-pad of the headpost directed the coil hotspot to the rat hindlimb motor cortex. A detachable coil guide of variable thickness can be used to further adjust coil positioning when necessary. The coil center could then be readily aligned to the hindlimb motor cortex by slightly moving the rat’s head along the medio-lateral direction until the left leg twitch was elicited.

### Habituation

Following surgery, the rats were allowed to recover before undergoing a 6-day habituation period. Habituation steps were performed to reduce stress during TMS administration. This involved holding the rats under the coil and playing recorded hdTBS sounds three times a day. Following this procedure, the rats showed significantly reduced stress, characterized by the absence of acute stress signs such as screaming, urination, defecation, and attempts to escape.

### Longitudinal hdTBS and I-O curve measurement

Following 6 days of habituation to the TMS environment and behavioral handling, we mapped the motor threshold and I-O curves on experimental Day 0. The rats were manually held with their hindlimb motor cortex region positioned under the hotspot of the coil (see Figure 1B). Their ears were manually blocked to minimize the effects of the acoustic noises from the TMS coil, which prevented the startle responses that would otherwise often be triggered if their ears were not blocked.

Motor threshold (MT), defined as the minimal TMS power to induce motor response in at least 50% of the trials, was determined by observing unilateral hindlimb motor twitches and MEP signal. I-O curves were obtained by delivering single pulses at 75% to 115% of the MT in 5% increment. A total of 8 pulses were delivered at each power level with an inter-pulse interval of 7 seconds. The resulting MEP signal was recorded; a separate channel recorded the trigger signal that indicated the timing of the TMS pulses.

Nine rats received hdTBS sessions from day 1 to day 5. hdTBS was identical to conventional iTBS except that each burst consisted of 6 pulses instead of 3 pulses, delivering a total of 1200 pulses in 190 seconds. TMS power level was set to 100% MT. A separate group of rats (N=7) received sham TMS. The experimental conditions in the sham group, including electrode/headpost implantation and habituation training, were identical to the TMS group except that the TMS power was set at 70% MT and the rat head was placed 4 cm beneath the TMS coil; the resulting electric field inside the rat brain was practically zero.

After 5 days of hdTBS administration, I-O curves were mapped on day 6 following identical procedures as described above.

### Longitudinal hdTBS and MRI Experiments

After determining that 1 hdTBS session/day for 5 days significantly enhanced cortical excitability as probed by I-O curves (see Results), we performed MRI experiments on another cohort of rats: N=8 for TMS, N=9 for sham. These rats underwent hdTBS or sham treatment using procedures as described above and received fMRI scans on days 0 and 6 under anesthesia. Because the MEP electrodes were not compatible with MRI, no electrodes were implanted on these animals, as a result, no I-O curve mapping was performed on these animals.

### MRI scan

A combination of isoflurane and dexmedetomidine was used to anesthetize rats during fMRI experiments, detailed procedure were described in [20,21]. MRI data were acquired with a Bruker Biospin 9.4T scanner running the Paravision 7.0 software platform (Bruker Medizintechnik, Karlsruhe, Germany). A volume quadrature transmitter coil (model: MT0381) was used for RF excitation, and a circular surface coil (model: MT0105-20) for MR signal reception. The decussation of the anterior commissure (approximately −0.36 mm from bregma) served as the fiducial landmark to standardize slice localization both within and across animals[22].

Structural images were collected using a Rapid Acquisition with Relaxation Enhancement sequence. Scan parameters: repetition time (TR) = 3100 ms, echo time (TE) = 36 ms, field of view (FOV) = 30 × 30 cm^2^, in-plane matrix size = 256 × 256, slice thickness = 0.6 mm, slice gap = 0.1 mm, slice number = 31.

Resting-state fMRI data were collected using gradient-echo echo-planner imaging (EPI) sequence developed in-house. fMRI data will be reported elsewhere. CBV-fMRI images were acquired after the resting scans, and they were collected using conventional multi-echo gradient-echo imaging (MGE) sequence. Scan parameters were: TR = 600 ms, TE = 2 ms, 10 echo images, 7 repetitions, slice number = 19, FOV = 30 ×30 mm^2^, and image size = 128 × 128. Immediately after the first two repetitions of data had been acquired, Monocrystalline Iron Oxide Nanoparticles (MION) agent was injected into the tail vein in 1 minute through an infusion pump (Feraheme, AMAG Pharmaceuticals Inc. Cambridge, MA, dose 15 mg/kg). The MION dose was chosen based on literature data[23,24] and our experiments[25], which was found to provide a good compromise between sensitivity and CBV weighting. MRI data was uninterrupted while MION infusion was taking place. The MGE scan lasted for 8 minutes and 58 seconds.

## Data Analysis

### MEP data analysis

I-O curve mapping was conducted at 9 TMS power levels (ranging from 75% to 115% MT). The MEP signal in rats features a latency of approximately 6 ms[14,26], the time stamps of the trigger signal was used to facilitate MEP identification. The peak-to-peak amplitudes of the MEP signal were quantified and averaged across trials for each power level. The averaged MEP of each rat at each power level was recorded as the output. The maximum amplitude of the baseline I-O curve was recorded, and the MEPs collected at each power level on different days were normalized based on the maximum baseline value, and log-transformed to ensure normal distribution[27]. For each group, the differences in MEP signal before and after the longitudinal modulation (Day 6 vs. Day 0) were compared using two-way repeated measures ANOVA with experimental Day (6 or 0) and TMS Power as the two factors, followed by *post hoc* paired t-test between power levels.

For both the real and sham hdTBS group, I-O curves on Day 0 (baseline) was subtracted from that on Day 6, the resulting I-O curve differences (delta-IO) reflect changes in cortical excitability due to 5 days of real or sham hdTBS, which were then subjected to two-way ANOVA (Stimulated Condition × TMS Power). We also computed the area under the curve (AUC) of the delta-IO for both the real and sham hdTBS groups. The AUC values were log-transformed to ensure normal distribution and were then subjected to two-sample t-tests for statistical comparison.

### CBV data analysis

fMRI data analyses were performed using two software packages: Analysis of Neuroimages (AFNI)[28] and FSL[29], along with custom scripts written in MATLAB. Initially, the CBV data underwent alignment with a template as follows: high-resolution RARE anatomical images were skull-stripped, linearly aligned to a template; the resulting transformation matrix was applied to the MGE images. Subsequently, relaxation time *R*2* was derived via a least-squares fitting of the MGE data, employing the following equation[30]:

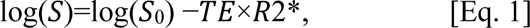

where *S* denotes MRI signal intensity, *S*_0_ signifies signal intensity at *TE*=0, and *TE* represents echo time.

This fitting was performed on a voxel-wise basis. To enhance the signal-to-noise ratio (SNR), the R2* values from the first two repetitions acquired before contrast agent injection (pre-MION) were averaged. Similarly, the R2* values from the last two repetitions (post-MION), obtained after contrast agent injection, were also averaged. Subsequently, the Δ*R*2* was computed by subtracting the pre-MION R2* from the post-MION R2*. Voxel-wise Δ*R*2* values are linearly correlated to corresponding CBV[31–34].

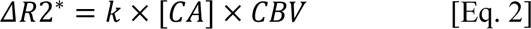

Where k is a constant, CA is the concentration of the contrast agent in blood.

CBV values at baseline (Day 0) and post-TMS (Day 6) in both the hdTBS treatment group and the sham group were computed in the same fashion. Changes in CBV values resulting from hdTBS or sham treatment were derived by calculating the differences in ΔR2* values between Day 0 and Day 6, which were subject to two-tailed t statistics, followed by corrections for multiple comparisons using the function 3dclustersim in AFNI (p<0.05, voxels in clusters>38).

## Results

hdTBS sessions were well tolerated by the animals. We observed no signs of abnormality in their daily behaviors, including eating, drinking or grooming, etc. Importantly, no signs of seizure events were observed in any of the animals during or after hdTBS administration.

As an example, Figure 1C shows raw MEP signals acquired on Day 0 from one representative rat. There were large amplitude artifacts immediately following the TMS pulses lasting for about 4 ms (indicated by the arrow), followed by multi-phasic MEP signal that featured a latency of about 6 ms (indicated by the oval circles).

### I-O curves at baseline and one day post-hdTBS modulation

Figure 3 summarizes I-O curve measurements on Day 0 (baseline) and Day 6 (one day post-TMS modulation). Two-way repeated ANOVA was done to evaluate the differences of the factors (Time x TMS Power). For the hdTBS group, there was significant TMS Power effect (*F*=16.97, degree of freedom (DOF) = 8, *p*<0.0001); there was also significant Time effect (Day 6 vs. Day 0) (*F*=9.63, DOF=1, *p*=0.0146). For the sham TMS group, there was significant TMS Power effect (*F*=21.80, DOF=8, *p*<0.0001), but no significant Time effect (*F*=2.07, DOF=1, *p*=0.20). Thus, MEP signal showed significant dependence on TMS power, as expected. Daily single-session of hdTBS for 5 days significantly shifted the I-O curve upward only in the TMS group (N=9, Figure 3A), but not in the sham group (N= 7, Figure 3B).

**Figure 2.**
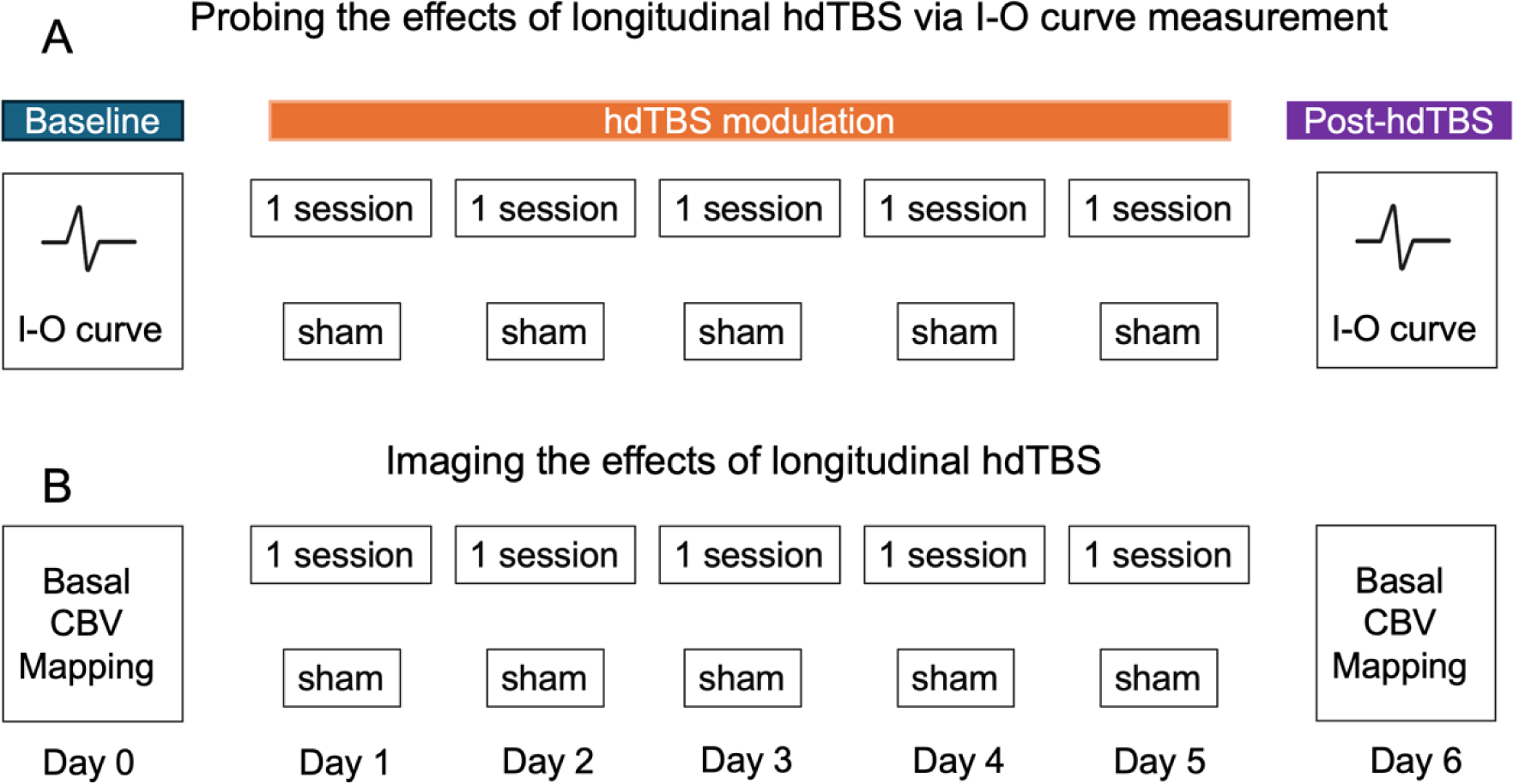
Illustration of experimental design on two separate cohorts of rats. (A): One cohort of rats received daily single-session of either real (N=8) or sham (N=7) hdTBS from day 1 to day 5. Input-output (I-O) curves were mapped on Day 0 and Day 6 to assess cortical excitability. (B): Another cohort of rats underwent identical hdTBS modulation procedures as in (A) (N= 8 for real hdTBS; N=9 for sham hdTBS), fMRI based on CBV mapping was performed on Day 0 and Day 6.

**Figure 3.**
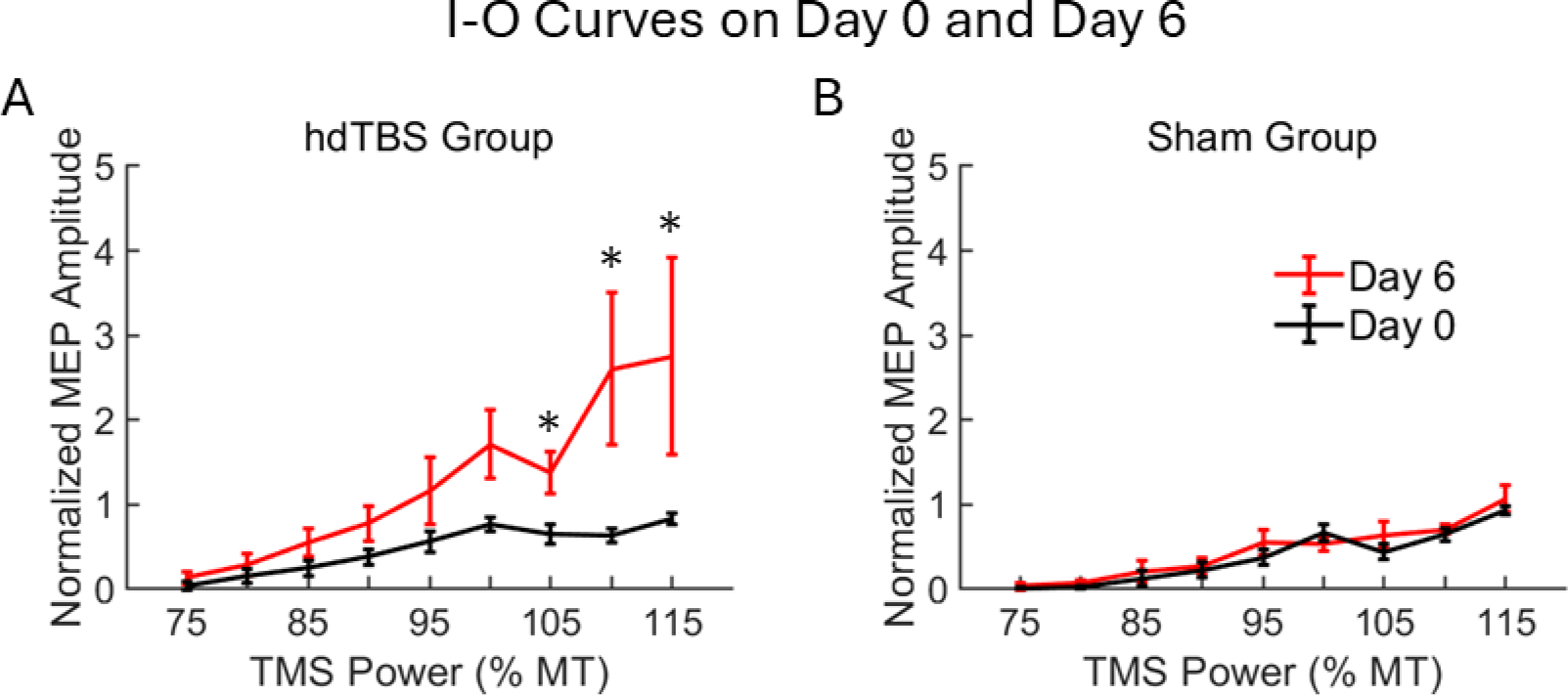
Input-output (I-O) curves were mapped at baseline (Day 0) and one day post-hdTBS modulation (Day 6). (A): Daily single-session of hdTBS for 5 days significantly shifted I-O curve upwards, while the I-O curves were indistinguishable in the sham TMS group (B).

We performed further analysis on the hdTBS group to determine the power levels that induced significant changes in MEP signal. *Post hoc* Results showed that at power levels of 105%, 110% and 115% MT, there were significant differences in MEP signal between Day 6 and Day 0 (*p*=0.0059, 0.0263 and 0.0419, respectively).

To further quantify alterations in I-O curves resulting from real or sham hdTBS, I-O curves at baseline (Day 0) were subtracted from that on Day 6, the resulting difference curves (delta-IO) in the real and sham hdTBS groups were subjected to two-way ANOVA (TMS Power × real/sham hdTBS Group), results are shown in Figure 4A. There was significant Group effect (*F*=11.84, DOF=1, *p*=0.0008). We also calculated the AUC values of delta-IO curves shown in Figure 4A from each animal, results are shown in Figure 4B. *Post hoc* study revealed AUC values in the hdTBS group was significantly higher than the sham group (*p*=0.03).

**Figure 4.**
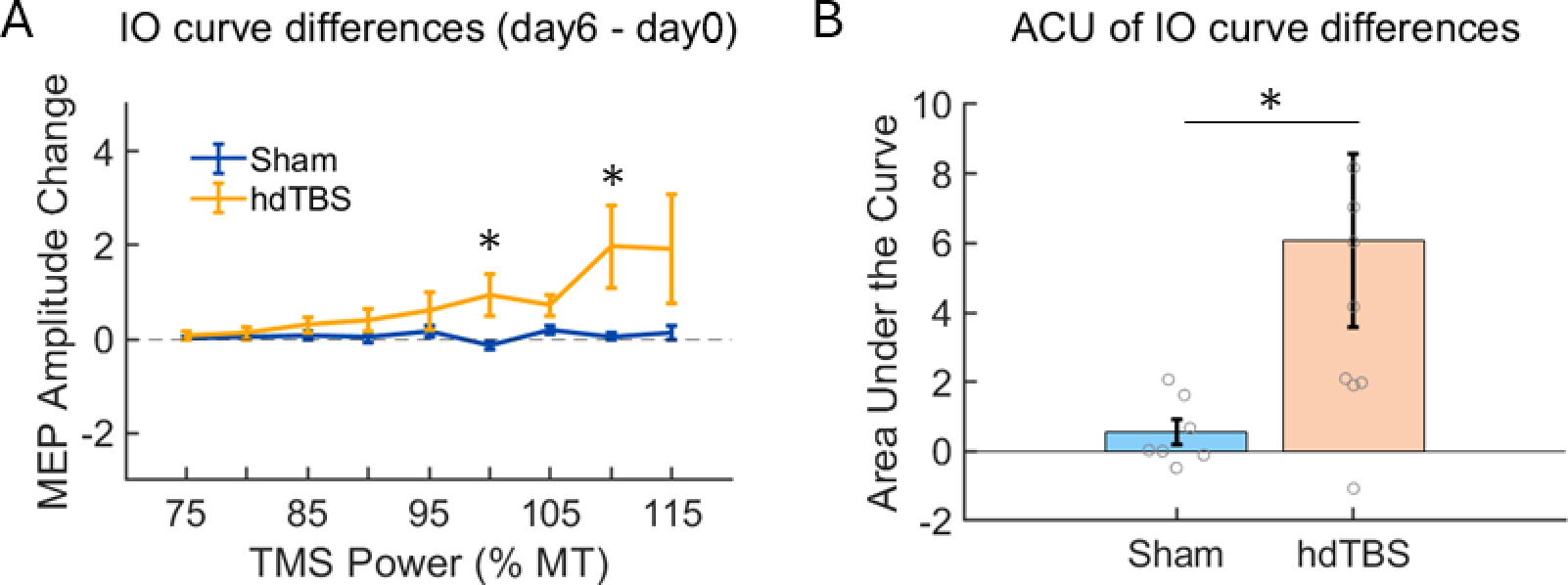
(A): I-O curve differences (delta-IO), calculated by subtracting I-O curves on Day 0 from that on Day 6, in the hdTBS (N=8) and sham group (N=7). (B): Area under the curve (AUC) of the delta-IO values in the real and sham hdTBS group (*, *p*=0.03).

### CBV mapping

Having discovered that daily single-hdTBS session for 5 days significantly shifted the I-O curves upward, indicating enhanced cortical excitability in the stimulated loci, we next investigated whether the functions of the brain regions interconnected with the stimulation loci were also altered by longitudinal TMS. To this end, we mapped basal CBV of the entire rat brain with the injection of intravascular contrast agent MION. As an example, Figure 5A shows a raw MGE image (TE=2 ms) before MION injection; Figure 5B shows the MRI signal intensities from one voxel with TEs ranging from 2 ms to 20 ms. Figure 5C-D shows the same imaging slice post-MION injection, along with a plot of the MRI signal across TEs. The TE-dependence of the MRI signal can be well described using a single exponential decay model (Figures 5B and 5D). As shown in this figure, and consistent with previous reports in rats and mice[23,24,30,32,33], MION injection drastically increased the relaxation rate (R2*) of the MRI signal.

**Figure 5.**
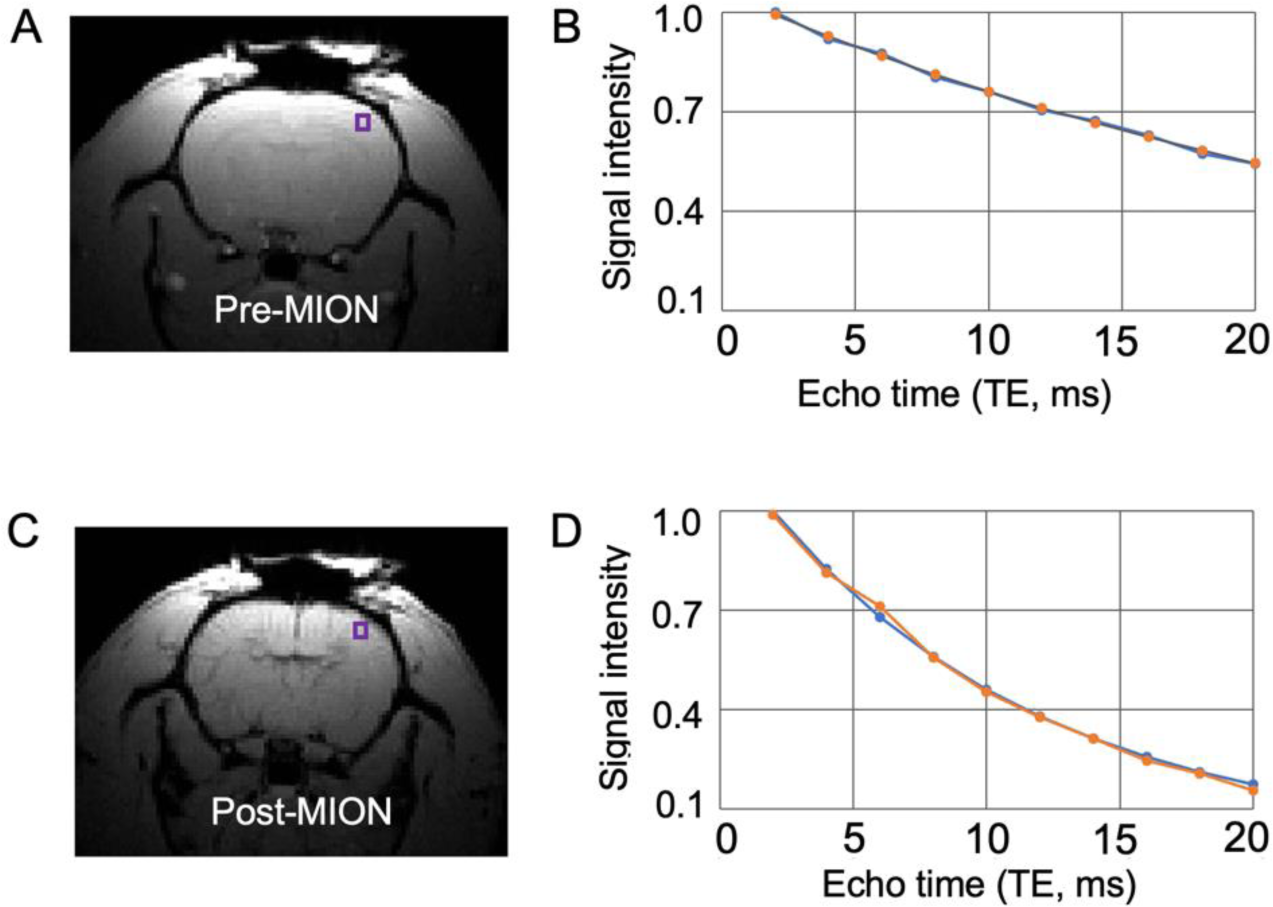
Visual comparison of MRI signal before and after contrast agent injection (MION, iron dose: 15 mg/kg). (A): Representative gradient echo image before MION injection, along with raw (orange) and fitted (blue) MRI signal over various echo times (TEs) shown in (B). (C-D) The same imaging slice after MION injection. MRI signal was chosen from the voxel indicated by the square boxes in A and C, respectively.

Changes in CBV values (ΔCBV) between Day 6 and Day 0 were calculated. Figure 6 shows voxel-wise statistical comparisons of ΔCBV values between the hdTBS group and the sham group. Significant differences were observed in the motor cortex (M1, the target site), somatosensory cortex of the hindlimb (S1HL), somatosensory cortex of the forelimb (S1FL), somatosensory barrel cortex (S1BF), caudate putamen (CP), amygdala, and visual cortex (V1, V2), showing a higher increase in CBV in the TMS group compared to the sham group. Figure 7 shows regional CBV values on Day 0 and Day 6 in both the TMS group and sham group.

**Figure 6.**
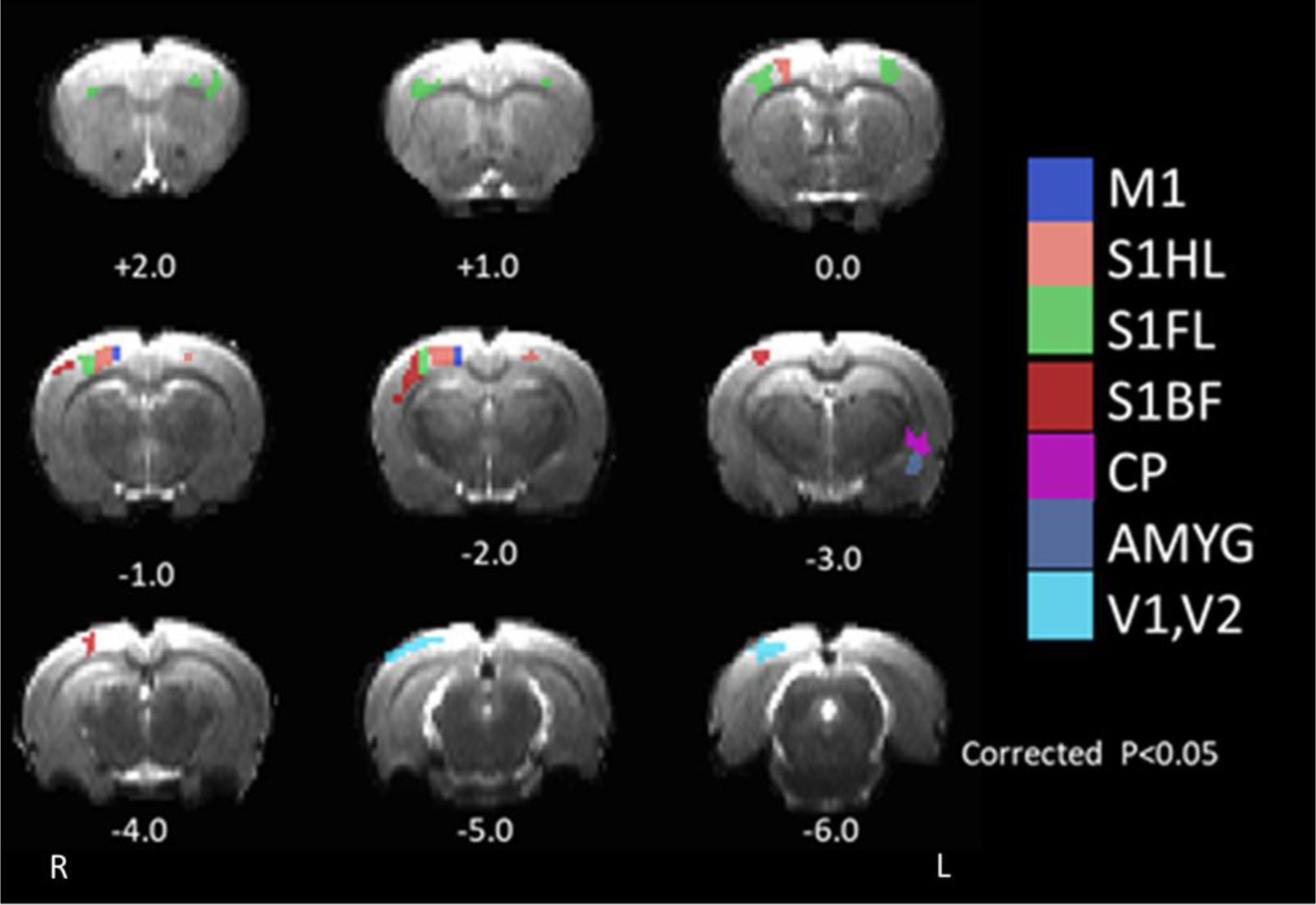
Statistical maps showing significant alterations in CBV values between the real and sham TMS group after daily single-session real (N=8) or sham (N=9) hdTBS for 5 days. *p*<0.05 after for multi-comparison correction.

**Figure 7.**
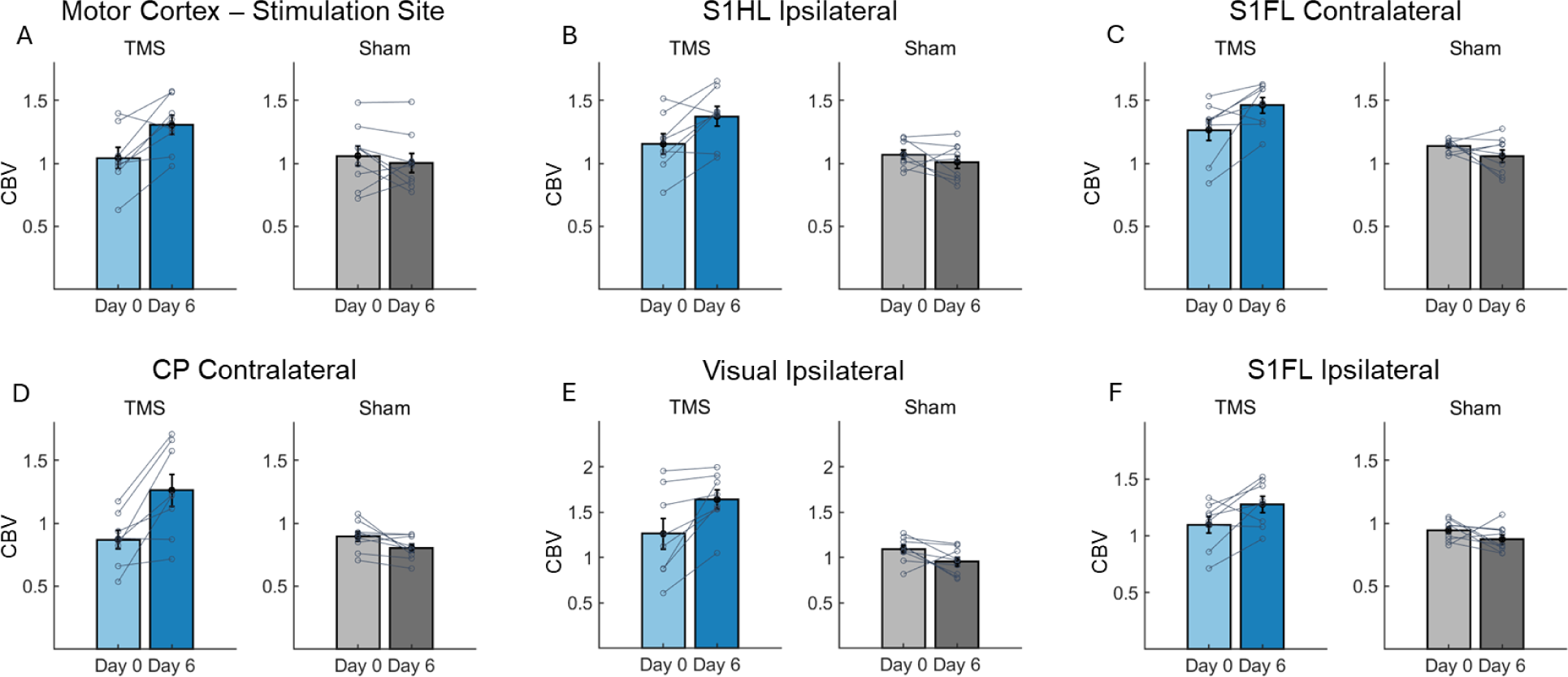
Bar plots of CBV changes in individual animals at baseline (Day 0) and one day post hdTBS modulation (Day 6).

## Discussion

In the present study, leveraging recent technological advancements including the focal TMS coil and hdTBS stimulator, combined with an awake rat model that allows for longitudinal MEP recording, we assessed neural plasticity of the rat motor cortex following hdTBS once daily for five days. The results reveal that this longitudinal TMS protocol significantly shifted the I-O curve upward, suggesting increased cortical excitability. Additionally, fMRI data demonstrated significant increases in CBV across several regions, including M1, S1HL, S1FL, S1BF, CP, and amygdala, indicating enhanced basal metabolism in these areas due to the longitudinal TMS. The imaging results aligned with previous acute stimulation effects in human and support the notion that TMS activation propagates across interconnected network beyond the stimulation loci[35–37].

This study utilizes established methodologies such as MEP readout, I-O curve assessment, and fMRI mapping, all of which are widely used in human research. Additionally, the sensorimotor cortex shows a high degree of conservation between rodents and humans, in contrast to high order cortical regions. These factors collectively strengthen the face validity of our rat model in back-translating TMS findings from humans to rodents.

### Neuroplasticity across cortical layers at the site of stimulation

The largest cluster exhibiting a significant increase in basal CBV was located ipsilateral to the stimulation site, centered at bregma -2 mm, corresponding to cortical representation of hindlimb motor cortex and adjacent sensory cortex[19]. This observation reaffirms the precise targeting ability of the coil. Furthermore, a small cluster in the contralateral sensorimotor region also exhibited an increase in CBV, likely influenced by interhemispheric callosal connections originating from the activated ipsilateral hemisphere.

fMRI based on CBV contrast can achieve laminar[30] and columnar resolution[38]. Our current CBV imaging data were acquired with an in-plane resolution of 234 μm, allowing us to explore the laminar profile of neuroplasticity induced by longitudinal TMS. Significant increases in CBV were observed in deep cortical layers, both ipsilateral and contralateral to the stimulation site (Figure 6). In a previous study mapping glucose uptake following acute TMS, we found significant increases in deep layers of the motor cortex[13]. Taken together, these findings indicate that TMS pulses activate deep cortical layers and that longitudinal TMS induces neuroplasticity in these regions.

The above experimental observation has strong implications in better understanding the mechanism of TMS action. Neuroanatomically, the mammalian neocortex exhibits distinct laminar profiles characterized by different types of neurons forming intricate circuits. Excitatory neurons, categorized by their projection targets, include intratelencephalic, pyramidal tract, and corticothalamic neurons[39]; Inhibitory neurons are distributed across laminae, forming excitatory-inhibitory local circuits [40,41]. Computational modeling[42] suggests that layer 5 pyramidal cells have the lowest activation thresholds, while layer 2/3 pyramidal cells and inhibitory basket cells are activated across most intensities; direct activation of layers 1 and 6 is considered unlikely. However, optical imaging of calcium signals suggests that TMS can directly activate layer 1 pyramidal neurons, while it may not directly activate dendrites of layer 5 neurons [43]. Our imaging data support the involvement of deep cortical layers resulting from suprathreshold TMS administration.

### Technical considerations

Anesthesia can introduce confounding variables in TMS studies, which can be particularly problematic in longitudinal applications where repeated anesthesia may overshadow or obscure the effects of repeated TMS. For instance, Ketamine, a commonly used anesthetic, is recognized for its potent antidepressant effects and is FDA-approved for treating major depression[44]. Gersner and colleagues compared TMS-induced neuroplasticity in awake rats versus rats under isoflurane anesthesia, revealing contrasting effects on neuroplasticity markers such as brain-derived neurotrophic factor (BDNF) levels and GluR1 subunit expression of AMPA receptors[45]. In the present study, TMS was administered exclusively in awake rats, avoiding the above confound, but our MRI imaging studies were performed with animals under the anesthesia. Anesthetics, depending on their molecular targets, could influence brain metabolism, affect neurovascular coupling pathways, and alter the arousal state[46]. The anesthesia protocol, comprising of low dose of dexmedetomidine and isoflurane, has been shown to preserve the synchrony of large scale brain networks [20,21], and has become the dominant method in the MRI field[47–49]. Since CBV mapping were conducted with animals under identical anesthesia conditions on Day 0 and Day 6, and both the real and sham hdTBS groups underwent the same anesthesia during imaging, it is not unreasonable to assume that the confounding effects from anesthesia are cancelled out in our statistical analysis.

Additionally, it would be interesting to explore the durability of the observed neuroplasticity. The current study mapped post-hdTBS CBV at only one time point (Day 6); future CBV mapping studies should be conducted at multiple time points to explore the time course of neuroplasticity induced by hdTBS.

## Acknowledgement

This work is supported in part by the Intramural Research Program of NIDA, NIH.

## Conflict of interest

**none.**

## Notes

### Competing Interest Statement

The authors have declared no competing interest.

### Summary of Updates

Nothing specific has changed. I added an author who contributed to the experiments, as we had mistakenly forgotten to include her.

## References

[1] Huang YZ, Edwards MJ, Rounis E, Bhatia KP, Rothwell JC. Theta burst stimulation of the human motor cortex. Neuron 2005;45:201–6. 10.1016/j.neuron.2004.12.033.

[2] Berlim MT, Eynde F van den, Tovar-Perdomo S, Daskalakis ZJ. Response, remission and drop-out rates following high-frequency repetitive transcranial magnetic stimulation (rTMS) for treating major depression: a systematic review and meta-analysis of randomized, double-blind and sham-controlled trials. Psychological Medicine 2014;44:225–39. 10.1017/S0033291713000512.

[3] Blumberger DM, Vila-Rodriguez F, Thorpe KE, Feffer K, Noda Y, Giacobbe P, et al. Effectiveness of theta burst versus high-frequency repetitive transcranial magnetic stimulation in patients with depression (THREE-D): a randomised non-inferiority trial. Lancet 2018;391:1683–92. 10.1016/s0140-6736(18)30295-2.

[4] Downar J, Blumberger DM, Daskalakis ZJ. Repetitive transcranial magnetic stimulation: an emerging treatment for medication-resistant depression. CMAJ 2016;188:1175–7. 10.1503/cmaj.151316.

[5] Eshel N, Keller CJ, Wu W, Jiang J, Mills-Finnerty C, Huemer J, et al. Global connectivity and local excitability changes underlie antidepressant effects of repetitive transcranial magnetic stimulation. Neuropsychopharmacology 2020;45:1018–25. 10.1038/s41386-020-0633-z.

[6] Cole EJ, Phillips AL, Bentzley BS, Stimpson KH, Nejad R, Barmak F, et al. Stanford Neuromodulation Therapy (SNT): A Double-Blind Randomized Controlled Trial. AJP 2022;179:132–41. 10.1176/appi.ajp.2021.20101429.

[7] Tiksnadi A, Murakami T, Wiratman W, Matsumoto H, Ugawa Y. Direct comparison of efficacy of the motor cortical plasticity induction and the interindividual variability between TBS and QPS. Brain Stimulation 2020;13:1824–33. 10.1016/j.brs.2020.10.014.

[8] Matsumoto H, Ugawa Y. Quadripulse stimulation (QPS). Exp Brain Res 2020;238:1619–25. 10.1007/s00221-020-05788-w.

[9] Wilson MT, Tang AD, Iyer K, McKee H, Waas J, Rodger J. The challenges of producing effective small coils for transcranial magnetic stimulation of mice. Biomed Phys Eng Express 2018;4:037002. 10.1088/2057-1976/aab525.

[10] Deng Z-D, Lisanby SH, Peterchev AV. Electric field depth–focality tradeoff in transcranial magnetic stimulation: simulation comparison of 50 coil designs. Brain Stimul 2013;6:1–13. 10.1016/j.brs.2012.02.005.

[11] Cohen LG, Roth BJ, Nilsson J, Dang N, Panizza M, Bandinelli S, et al. Effects of coil design on delivery of focal magnetic stimulation. Technical considerations. Electroencephalogr Clin Neurophysiol 1990;75:350–7.

[12] Meng Q, Jing L, Badjo JP, Du X, Hong E, Yang Y, et al. A novel transcranial magnetic stimulator for focal stimulation of rodent brain. Brain Stimul 2018;11:663–665. doi: 10.1016/j.brs.2018.02.018. Epub 2018 Mar 1.

[13] Cermak S, Meng Q, Peng K, Baldwin S, Mejías-Aponte CA, Yang Y, et al. Focal transcranial magnetic stimulation in awake rats: enhanced glucose uptake in deep cortical layers. J Neurosci Methods 2020;339:108709. 10.1016/j.jneumeth.2020.108709.

[14] Meng Q, Nguyen H, Vrana A, Baldwin S, Li CQ, Giles A, et al. A high-density theta burst paradigm enhances the aftereffects of transcranial magnetic stimulation: Evidence from focal stimulation of rat motor cortex. Brain Stimul 2022;15:833–42. 10.1016/j.brs.2022.05.017.

[15] GRUBB RL, RAICHLE ME, EICHLING JO, TER-POGOSSIAN MM. The Effects of Changes in PaCO2 Cerebral Blood Volume, Blood Flow, and Vascular Mean Transit Time. Stroke 1974;5:630–9. 10.1161/01.STR.5.5.630.

[16] Raichle ME. A brief history of human brain mapping. Trends Neurosci 2009;32:118–26. 10.1016/j.tins.2008.11.001.

[17] Pascual-Leone A, Valls-Solé J, Wassermann EM, Hallett M. Responses to rapid-rate transcranial magnetic stimulation of the human motor cortex. Brain 1994;117 (Pt 4):847– 58.

[18] Chen R, Classen J, Gerloff C, Celnik P, Wassermann EM, Hallett M, et al. Depression of motor cortex excitability by low-frequency transcranial magnetic stimulation. Neurology 1997;48:1398–403. 10.1212/wnl.48.5.1398.

[19] Seong HY, Cho JY, Choi BS, Min JK, Kim YH, Roh SW, et al. Analysis on Bilateral Hindlimb Mapping in Motor Cortex of the Rat by an Intracortical Microstimulation Method. J Korean Med Sci 2014;29:587–92. 10.3346/jkms.2014.29.4.587.

[20] Brynildsen JK, Hsu LM, Ross TJ, Stein EA, Yang Y, Lu H. Physiological characterization of a robust survival rodent fMRI method. Magn Reson Imaging 2016. 10.1016/j.mri.2016.08.010.

[21] Lu H, Zou Q, Gu H, Raichle ME, Stein EA, Yang Y. Rat brains also have a default mode network. Proceedings of the National Academy of Sciences 2012;109:3979–84. 10.1073/pnas.1200506109.

[22] Lu H, Xi ZX, Gitajn L, Rea W, Yang Y, Stein EA. Cocaine-induced brain activation detected by dynamic manganese-enhanced magnetic resonance imaging (MEMRI). Proc Natl Acad Sci U S A 2007;104:2489–94. 10.1073/pnas.0606983104.

[23] Pohlmann A, Karczewski P, Ku M-C, Dieringer B, Waiczies H, Wisbrun N, et al. Cerebral blood volume estimation by ferumoxytol-enhanced steady-state MRI at 9.4 T reveals microvascular impact of α1-adrenergic receptor antibodies. NMR in Biomedicine 2014;27:1085–93. 10.1002/nbm.3160.

[24] Moreno H, Hua F, Brown T, Small S. Longitudinal mapping of mouse cerebral blood volume with MRI. NMR Biomed 2006;19:535–43. 10.1002/nbm.1022.

[25] Lu H, Scholl CA, Zuo Y, Stein EA, Yang Y. Quantifying the blood oxygenation level dependent effect in cerebral blood volume-weighted functional MRI at 9.4T. Magn Reson Med 2007;58:616–21. 10.1002/mrm.21354.

[26] Rotenberg A, Muller PA, Vahabzadeh-Hagh AM, Navarro X, López-Vales R, Pascual-Leone A, et al. Lateralization of forelimb motor evoked potentials by transcranial magnetic stimulation in rats. Clin Neurophysiol 2010;121:104–8. 10.1016/j.clinph.2009.09.008.

[27] Wassermann EM. Variation in the response to transcranial magnetic brain stimulation in the general population. Clin Neurophysiol 2002;113:1165–71. 10.1016/s1388-2457(02)00144-x.

[28] Cox R, Hyde J. Software tools for analysis and visualization of fMRI data. NMR Biomed 1997;10:171–8. 10.1002/(SICI)1099-1492(199706/08)10:4/5<171::AID-NBM453>3.0.CO;2-L.

[29] Smith SM, Jenkinson M, Woolrich MW, Beckmann CF, Behrens TE, Johansen-Berg H, et al. Advances in functional and structural MR image analysis and implementation as FSL. Neuroimage 2004;23 Suppl 1:S208–19. 10.1016/j.neuroimage.2004.07.051.

[30] Lu H, Patel S, Luo F, Li SJ, Hillard CJ, Ward BD, et al. Spatial correlations of laminar BOLD and CBV responses to rat whisker stimulation with neuronal activity localized by Fos expression. Magn Reson Med 2004;52:1060–8. 10.1002/mrm.20265.

[31] Mandeville JB, Jenkins BG, Chen YCI, Choi JK, Kim YR, Belen D, et al. Exogenous contrast agent improves sensitivity of gradient-echo functional magnetic resonance imaging at 9.4 T. Magnetic Resonance in Medicine 2004;52:1272–81. 10.1002/mrm.20278.

[32] Wu EX, Wong KK, Andrassy M, Tang H. High-resolution in vivo CBV mapping with MRI in wild-type mice. Magnetic Resonance in Medicine 2003;49:765–70. 10.1002/mrm.10425.

[33] Mandeville JB, Marota JJ, Kosofsky BE, Keltner JR, Weissleder R, Rosen BR, et al. Dynamic functional imaging of relative cerebral blood volume during rat forepaw stimulation. Magn Reson Med 1998;39:615–24.

[34] Boxerman JL, Hamberg LM, Rosen BR, Weisskoff RM. Mr contrast due to intravascular magnetic susceptibility perturbations. Magnetic Resonance in Medicine 1995;34:555–66. 10.1002/mrm.1910340412.

[35] Bestmann S, Baudewig J, Siebner HR, Rothwell JC, Frahm J. Subthreshold high-frequency TMS of human primary motor cortex modulates interconnected frontal motor areas as detected by interleaved fMRI-TMS. Neuroimage 2003;20:1685–96. 10.1016/j.neuroimage.2003.07.028.

[36] Hampson M, Hoffman RE. Transcranial magnetic stimulation and connectivity mapping: tools for studying the neural bases of brain disorders. Front Syst Neurosci 2010;4:40. 10.3389/fnsys.2010.00040.

[37] Rossini PM, Burke D, Chen R, Cohen LG, Daskalakis Z, Di Iorio R, et al. Non-invasive electrical and magnetic stimulation of the brain, spinal cord, roots and peripheral nerves: Basic principles and procedures for routine clinical and research application. An updated report from an I.F.C.N. Committee. Clin Neurophysiol 2015;126:1071–107. 10.1016/j.clinph.2015.02.001.

[38] Fukuda M, Moon CH, Wang P, Kim SG. Mapping iso-orientation columns by contrast agent-enhanced functional magnetic resonance imaging: Reproducibility, specificity, and evaluation by optical imaging of intrinsic signal. Journal of Neuroscience 2006;26:11821–32. 10.1523/jneurosci.3098-06.2006.

[39] Harris KD, Shepherd GMG. The neocortical circuit: themes and variations. Nat Neurosci 2015;18:170–81. 10.1038/nn.3917.

[40] Douglas RJ, Martin KA. Neuronal circuits of the neocortex. Annu Rev Neurosci 2004;27:419–51. 10.1146/annurev.neuro.27.070203.144152.

[41] Douglas RJ, Martin KA. Inhibition in cortical circuits. Curr Biol 2009;19:R398–402. 10.1016/j.cub.2009.03.003.

[42] Aberra AS, Peterchev AV, Grill WM. Biophysically realistic neuron models for simulation of cortical stimulation. J Neural Eng 2018;15:066023. 10.1088/1741-2552/aadbb1.

[43] Murphy SC, Palmer LM, Nyffeler T, Müri RM, Larkum ME. Transcranial magnetic stimulation (TMS) inhibits cortical dendrites. Elife 2016;5. 10.7554/eLife.13598.

[44] Yavi M, Lee H, Henter ID, Park LT, Zarate CA. Ketamine treatment for depression: a review. Discov Ment Health 2022;2:9. 10.1007/s44192-022-00012-3.

[45] Gersner R, Kravetz E, Feil J, Pell G, Zangen A. Long-term effects of repetitive transcranial magnetic stimulation on markers for neuroplasticity: differential outcomes in anesthetized and awake animals. J Neurosci 2011;31:7521–6. 10.1523/JNEUROSCI.6751-10.2011.

[46] Brown EN, Purdon PL, Van Dort CJ. General anesthesia and altered states of arousal: A systems neuroscience analysis. In: Hyman SE, Jessell TM, Shatz CJ, Stevens CF, Zoghbi HY, editors. Annual Review of Neuroscience, Vol 34, vol. 34, 2011, p. 601–28. 10.1146/annurev-neuro-060909-153200.

[47] Grandjean J, Desrosiers-Gregoire G, Anckaerts C, Angeles-Valdez D, Ayad F, Barrière DA, et al. A consensus protocol for functional connectivity analysis in the rat brain. Nat Neurosci 2023;26:673–81. 10.1038/s41593-023-01286-8.

[48] Ortiz-Rios M, Sirmpilatze N, König J, Boreitus S. An anesthetic protocol for preserving functional network structure in the marmoset monkey brain. Imaging Neuroscience 2024;2:1–23. 10.1162/imag_a_00230.

[49] Kint LT, Seewoo BJ, Hyndman TH, Clarke MW, Edwards SH, Rodger J, et al. The Pharmacokinetics of Medetomidine Administered Subcutaneously during Isoflurane Anaesthesia in Sprague-Dawley Rats. Animals (Basel) 2020;10. 10.3390/ani10061050.

